# Intracellular K^+^ limits T cell exhaustion and preserves antitumor function

**DOI:** 10.1101/2023.09.13.556997

**Authors:** Camille Collier, Kelly Wucherer, Matthew McWhorter, Chelsea Jenkins, Alexandra Bartlett, Rahul Roychoudhuri, Robert Eil

## Abstract

The cancer-killing activity of T cells is often compromised within tumors, allowing disease progression. We previously found that intratumoral elevations in extracellular K^+^ related to ongoing cell death constrained CD8^+^ T cell Akt-mTOR signaling and effector function (1,2). To alleviate K^+^ mediated T cell suppression, we pursued genetic means to lower intracellular K^+^.

Transcriptomic analysis of CD8^+^ T cells demonstrated the Na^+^/K^+^ ATPase to be robustly and dynamically expressed. CRISPR-Cas9 mediated deletion of the catalytic alpha subunit of the Na^+^/K^+^ ATPase lowered intracellular K^+^ but produced tonic hyperactivity in multiple signal transduction cascades along with the acquisition of co-inhibitory receptors and terminal differentiation in mouse and human CD8^+^ T cells. Mechanistically, Na^+^/K^+^ ATPase disruption led to ROS accumulation due to depletion of intracellular K^+^ in T cells. Antioxidant treatment or high K^+^ media prevented *Atp1a1* deficient T cells from exhausted T (T_Ex_) cell formation. Consistent with transcriptional and proteomic data suggesting a T_Ex_ cell phenotype, T cells lacking *Atp1a1* had compromised persistence and antitumor activity in a syngeneic model of orthotopic murine melanoma. Translational application of these findings will include efforts to lower intracellular K^+^ while limiting ROS accumulation within tumor specific T cells.

**Synopsis:** High extracellular K^+^ (↑[K^+^]_e_) is found within tumors and suppresses T cell effector function. Collier et al. find that deletion of the Na^+^/K^+^ ATPase in T cells lowers intracellular K^+^ and promotes ROS accumulation, tonic signal transduction and T cell exhaustion owing to ROS accumulation. Engineering T cell ion transport is an important consideration for cancer immunotherapy.

## Introduction/ background

The past 20 years of translational oncology provide a framework in which the ability to eradicate cancer has increased in direct relation to our understanding of T cell biology (3). While T cells infiltrate cancers, only a small proportion of patients that receive immunotherapy realize durable regression of disease owing to tumor-induced suppression of T cell function (4–6). Following T cell receptor (TCR) engagement, naïve T cells (T_N_) progressively differentiate into stem cell memory (T_SCM_), central memory (T_CM_), effector memory (T_EM_), and terminally differentiated exhausted T cells (T_Ex_). As T cells differentiate along this continuum, transcriptional, epigenetic, and metabolic changes drive the acquisition of effector functions (IFN-γ production, cytolysis) in T_EM_ cells, along with the coincident loss of stem-like behaviors (multipotency, persistence, and self-renewal) seen in T_SCM_ and T_CM_ cells (7). Within cancers, repetitive TCR stimulation drives T_EM_ cells to differentiate into T_Ex_ cells with compromised effector functions and loss of stem-cell like behavior. In addition to T_Ex_ cell hyporesponsiveness, direct suppression of TCR signaling (i.e. PD-L1, CTLA-4) can drive T cell intratumoral dysfunction (3,4). We previously reported that elevated extracellular potassium (↑[K^+^]_e_) within tumors directly suppresses TCR induced signal transduction and effector function (1,2,8). In the current work, we aimed to imbue tumor specific T cells with resistance to ↑[K^+^]_e_ by reprogramming K^+^ transport to result in a lower intracellular K^+^.

To understand the underpinnings of T cell K^+^ physiology, we performed whole-transcriptomic analysis of T cells in varied states of activation and differentiation. While our efforts and those of others have focused on the role of the K^+^ channels K_v_1.3 and K_Ca_3.1, we also found that transcripts encoding a number of other K^+^ channels and pumps were differentially abundant across T cell populations (9). We elected to focus on the Na^+^/K^+^ ATPase due owing to its established contribution to intracellular [K^+^] and its robust expression. At the expense of a significant portion of total cellular ATP, the Na^+^/K^+^ ATPase exports three sodium ions and imports two potassium ions. The result is a cytoplasm that is hyperkalemic and hyponatremic with an electrical potential (i.e. *V*_m_) of approximately -60 mV compared to the extracellular space (10). Here, we found that, despite disrupting the transmembrane electrical gradient, lowering intracellular [K^+^] and impairing Ca^2+^ influx, genetic disruption of the Na^+^-K^+^-ATPase produced spontaneous activation within TCR and Akt-mTOR signaling cascades. This resulted in terminal differentiation and T_Ex_ cell formation, mirroring other instances of tonic T cell signaling. Mechanistically, we found T cells without an intact *Atp1a1* locus to be in a state of stress response, with elevated levels of reactive oxygen species (ROS). Restoration of intracellular K^+^ levels or supplementation with the antioxidant N-acetylcysteine (NAC) prevented ROS accumulation, terminal differentiation, and T_Ex_ cell formation. Consistent with their T_Ex_ cell surface phenotype, tumor specific T cells lacking *Atp1a1* had compromised *in vivo* persistence and antitumor activity compared to control treated cells in a mouse model of orthotopic melanoma. These findings indicate that low intracellular [K^+^] in T cells produces unregulated signal transduction activity in multiple cascades, including the Akt-mTOR pathway, resulting in T cell exhaustion. Future efforts to render T cells resistant tothe stressors of the tumor microenvironment will require balancing K^+^ concentration with ROS abundance, activation state and effector function.. Broadly, a deeper understanding of T cell ion transport will advance cancer immunotherapy by informing the development and application of pharmacologic and genetic interventions that preserve T cell stem-cell like behaviors while also augmenting intratumoral T cell effector function.

## Results and Discussion

### Cell intrinsic regulators of T cell ion transport

We previously found that cancer cell death results in accumulation of K^+^ within the extracellular space of tumors, constraining mouse and human T cell activation (1,11). Here, we find that extracellular K^+^ characteristic of tumors (↑[K^+^]_e_) significantly suppresses cytokine production in response to TCR ligation in CD8^+^ T cells (**Figure 1A**). Additionally, we found that ↑[K^+^]_e_ enacted T cell suppression via elevations to intracellular K^+^ ([K^+^]_i_), as demonstrated by increased fluorescence of the K^+^-sensitive dye APG-4 (**Figure 1B**). Use of the ionophore gramicidin lowered [K^+^]_i_ and restored cytokine production.

**Figure 1.**
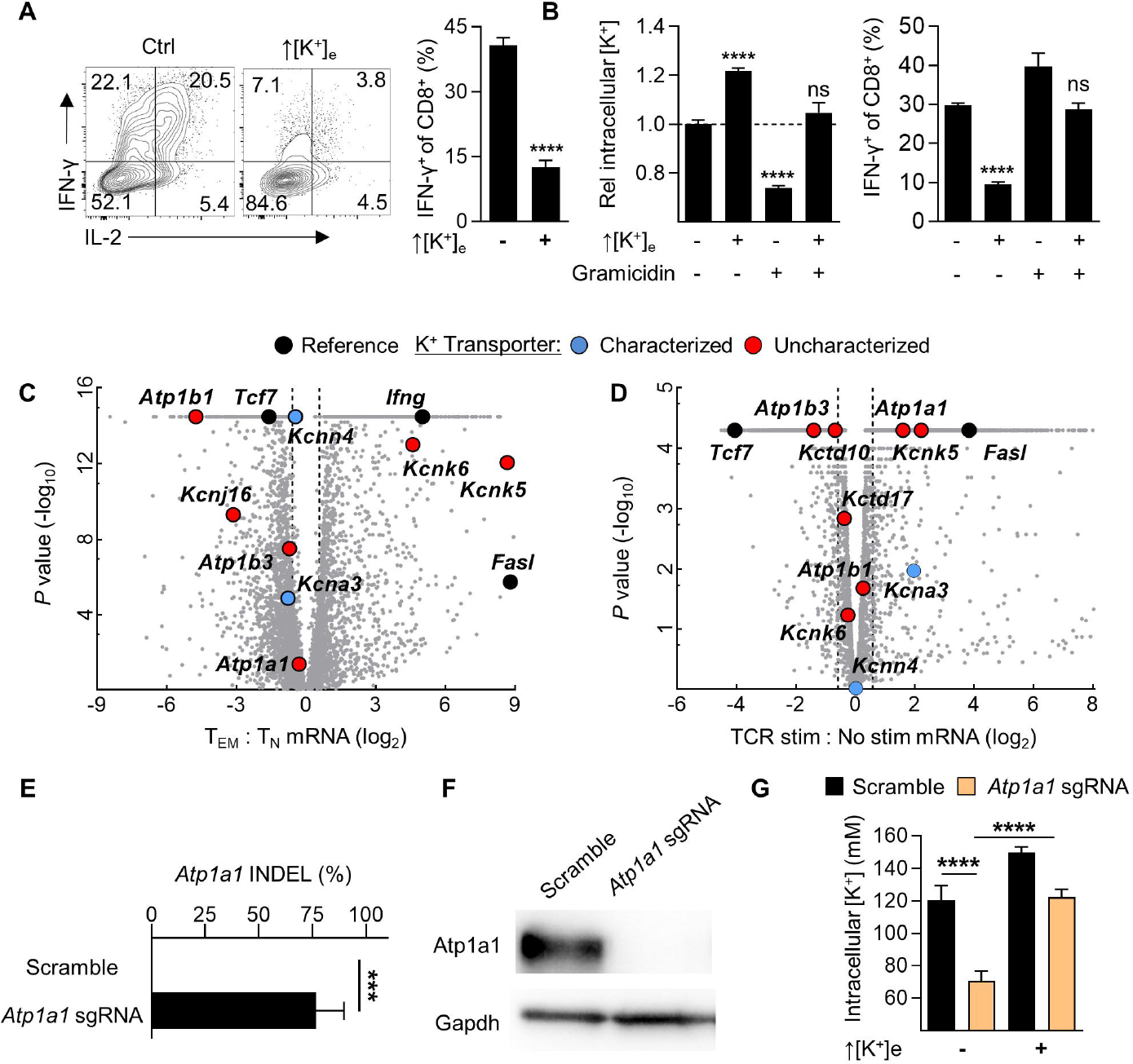
Figure 1. Elevated extracellular K^+^ in tumors suppresses T cell function via increasing intracellular levels. **(A)** Representative flow cytometry plots and summary bar graphs of activated mouse CD8^+^ of T cells acutely restimulated for 5 hours with immobilized aCD3&28 as indicated after being expanded in Ctrl conditions. **(B)** Summary flow cytometry bar graphs of CD8+ T cells exposed the indicated conditions (↑[K^+^]_e_ = 40 mM; Gramicidin = 1.5µM). Intracellular [K^+^] is captured by the K^+^-sensitive dye APG-4, whose fluorescence is displayed in ratio to the vehicle condition. **(C)** Volcano plot depicting comparative expression of loci amongst CD8^+^ OT-I natïve (T_N_) and effector memory (T_EM_) T cells or **(D)** Comparing global transcriptomic changes induced by TCR ligation in effector T cells. **(E)** INOEL quantification following CRISPR-Cas9 mediated disruption of the *Atp1a1* locus in CD8^+^ and **(F)** quantification of protein abundance by immunoblot of the same cells. **(G)** Summary bar graphs quantifying [K^+^]_i_ of CD8^+^ T cells following CRISPR-Cas9 mediated disruption (Scramble, sgRNA *Atp1a1)*, as indicated by the K^+^-sensitive, fluorometric dye APG-4, captured via flow cytometry (↑[K^+^]_e_ = 60 mM). Error bars represent standard deviation. *P < 0.05; ***P < 0.005; ****P < 0.001, 2-tailed Student’s t tests **(A-E,G)**.

T cell K^+^ transport has not been comprehensively characterized. To obtain a holistic understanding of K^+^ transporters in T cells, we FACS isolated naïve (T_N_) CD44^-^CD62L^+^ CD8^+^ T cells from OT-I TCR transgenic *Rag2*^-/-^CD45.2^+^ mice and transferred them to CD45.1^+^ hosts in combination with ovalbumin expressing vaccinia virus (VV-OVA). We then purified T_N_ (CD44^-^CD62L^+^) and T_EM_ (CD44^-^CD62L^-^) populations from the recipients and performed whole-transcriptomic analysis (**Figure 1C**). Similarly, we activated CD8^+^ T cells for 2 hours via their TCR to induce effector function and assessed global transcriptional changes (**Figure 1D**).

We found high levels of transcripts encoding the channels K_v_1.3 (*Kcna3*) and K_Ca_3.1 (*Kcnn4*), as expected. However, multiple other voltage gated, ligand gated, two-pore, and inwardly-rectifying channels along with alpha and beta subunits of the Na^+^/K^+^ ATPase were differentially expressed across CD8^+^ subset and state. Drawing from these findings, we elected to interrogate the role of the Na^+^/K^+^ ATPase in CD8^+^ T cells owing to its abundance and dynamism in CD8^+^ T cells, along with its importance in other cell types that maintain high intracellular [K^+^] (12).

### The Na^+^-K^+^-ATPase is required for T cell quiescence

The Na^+^/K^+^ ATPase is a heterodimeric enzyme consisting of catalytic (alpha) and regulatory (beta) subunits (12). Since both *Atp1b1* and *Atp1b3* isoforms of the β subunit are present in CD8^+^ T cells, we targeted the nonredundant α isoform, encoded by the *Atp1a1* locus, for genetic disruption (**Figure 1E**,**F**). Using the fluorometric K^+^ dye APG-4, we found that CD8^+^ T cells lacking the Na^+^/K^+^ ATPase harbored an intracellular [K^+^]_i_ of 70.6±6mM vs 130.8±9mM in control treated cells (**Figure 1G**). In addition to setting a high [K^+^]_i_, the Na^+^/K^+^ ATPase maintains negative resting membrane potential (*V*_m_) relative to the extracellular space (12). We used the voltage sensitive membrane permeable dye DiSBAC_2_(3), also known as oxonol, to comparatively assess membrane potential between CD8^+^ T cell populations. As expected, T cells deficient for *Atp1a1* maintained a depolarized (closer to zero) cellular *V*_m_ compared to scramble treated controls. CD8^+^ T cells transduced with retroviral particles encoding K^+^ channels that enforce a more negative membrane potential, i.e. hyperpolarization (K_ir_2.1 over expression), or increase the membrane potential closer to zero, depolarization (dominant negative K_v_1.3), are included for comparison and assay validation (**Figure 2A**). We also found *Atp1a1* disruption to compromise Ca^2+^ influx following both TCR and ionomycin induced store operated calcium entry (**Figure 2B, C**). Despite decreased TCR induced Ca^2+^ influx, CD8^+^ T cells lacking the Na^+^-K^+^-ATPase had abundant phosphorylation within a number of signal transduction pathways. Most notably, CD3ζ sustained high levels of phosphorylation along with Akt-mTOR and MAPK/ Erk pathway members (**Figure 2D**). Consistent with increased activation within multiple signaling cascades, we found CD8^+^ T cells targeted for *Atp1a1* disruption maintained high levels of activation and exhaustion markers such as PD-1, Tim-3, Fas, and CD69 with a relative loss of the stemness associated proteins such as Tcf7 and CD62L (**Figure 2E,F**). These attributes ultimately diminished effector function in *Atp1a1-*deficient cells (**Figure 2G**). The prominent PD-1^+^Tim-3^+^ expression suggests accelerated differentiation towards a terminal T_Ex_ cell state. Collectively, these data indicate that, despite promoting TCR induced Ca^2+^ influx and subsequent T cell activation, the Na^+^/K^+^ ATPase is required to constrain tonic signal transduction that otherwise disrupts T cell quiescence and promotes terminal differentiation.

**Figure 2.**
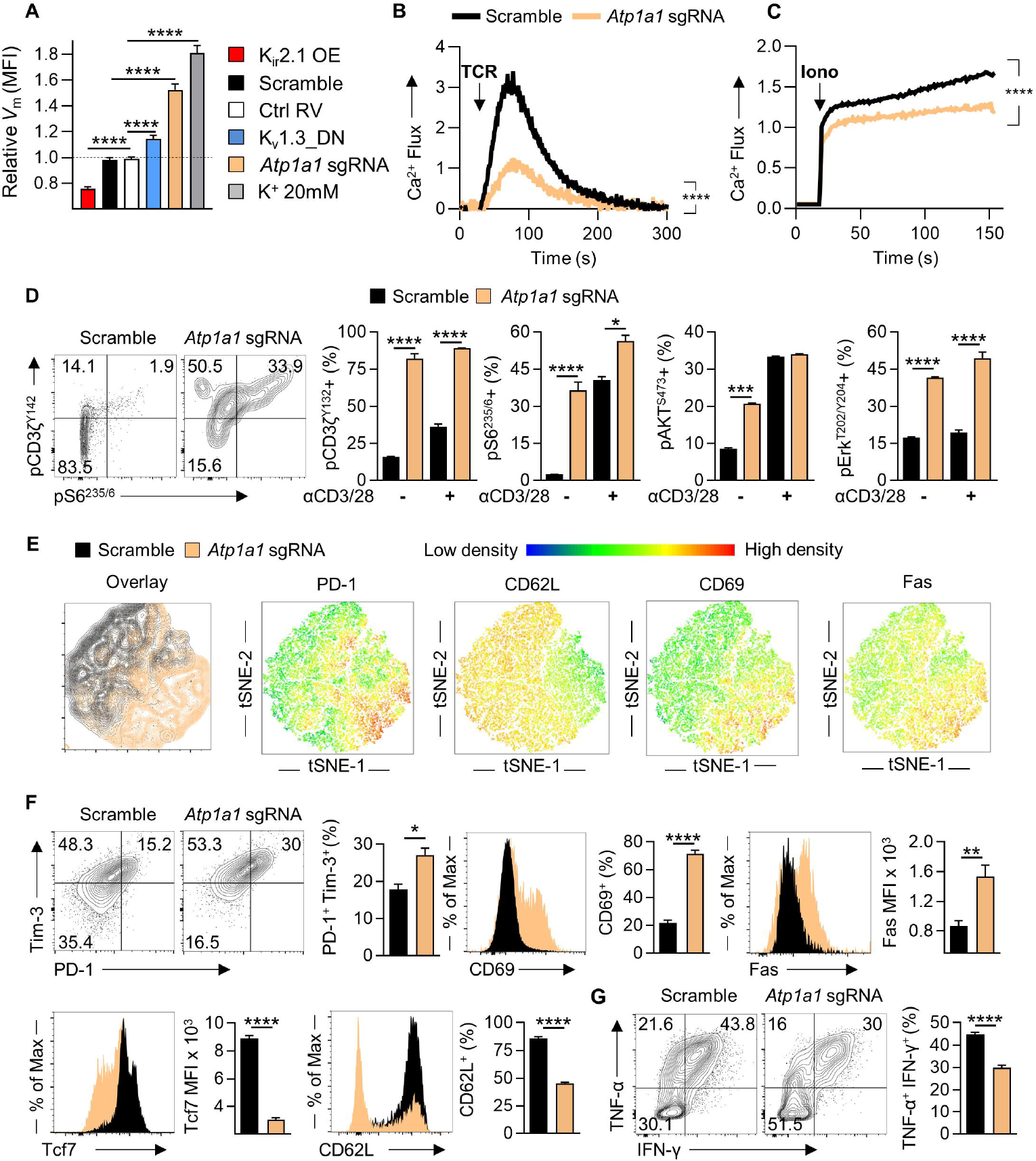
The Na^+^/K^+^ ATPase restrains tonic signaling and CD8^+^ T cell dysfunction. **(A)** Summary bar graphs quantifying fluorescence of the voltage sensitive indicator DiSBAC_2_3, reflecting the transmembrane electrochemical gradient (V_m_) of CD8^+^ T cells in standard RPMI 1640 (Vehicle) or isotonic hyperkalemic conditions (**K**^+^ 20mM) captured by flow cytometry following CRISPR-Cas9 mediated disruption (Scramble, sgRNA *Atp1a1)* or transduction with retroviral particles. OE, over-expression; DN, Dominant negative. **(B,C)** Summary quantification of cytoplasmic Ca^2+^ as interval Fluo-3/ FuraRed fluorescence following TCR cross-linking **(B)** or ionomycin induced store-operated calcium flux **(C)** as indicated captured by flow cytometry. lono, lonomycin. Average of 3 technical replicates depicted, representative of 2 independent experiments. (**D**) Representative phosflow cytometry plots and summary quantification of CD8^+^ T cell populations in the or absence of soluble TCR cross-linking stimulation for five minutes. (**E**) Concatenated single-cell overlay and colorimetric density tSNE based depiction of the indicated proteins captured by flow cytometry, **(F)** Representative flow cytometry plots and summary quantification for the indicated markers in *ex vivo* expanded CD8^+^ T cells. **(G)** Representative and summary quantification of cytokines following acute TCR re-stimulation of *ex vivo* expanded CD8^+^ T cells. Error bars represent standard deviation. *P < 0.05; ***P < 0.005; ****P < 0.001, 2-tailed Student’s t tests **(D,F,G)**. ****P < 0.001 for two-way ANOVA **(B,C)**.

### The Na^+^-K^+^-ATPase supports ROS homeostasis to restrain CD8^+^ T cell terminal differentiation

To better understand how the Na^+^/K^+^ ATPase controls T cell behavior, we examined how *Atp1a1* impacts T cell differentiation and function using orthogonal approaches. First, we performed whole-transcriptomic analysis of C57BL/6 CD8^+^ splenocytes four days after CRISPR-Cas9 mediated disruption of *Atp1a1*. Consistent with our flow cytometry readouts, we found *Atp1a1* disruption promoted effector T cell programs, stress response pathways, and anabolic metabolism with a coincident loss of stemness associated programs (**Figure 3A,B**). Gene set enrichment analysis for expression profiles increased following *Atp1a1* disruption identified signatures canonical for stress response, effector T cell programs, Mtorc1 and ROS metabolism among others (**Figure 3C,D**).

**Figure 3.**
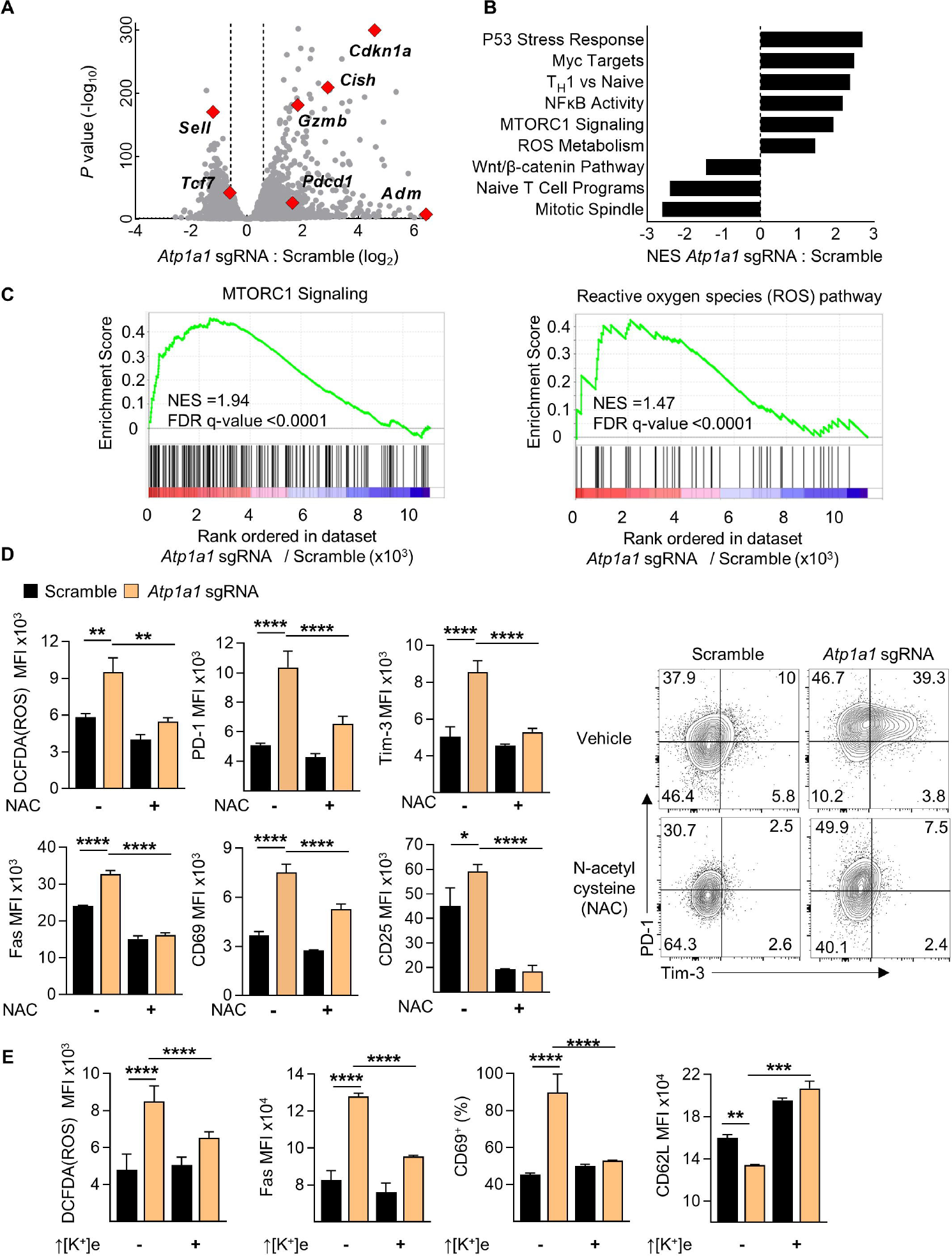
The Na^+^/K^+^ATPase limits T cell dysfunction via maintaining intracellular [K^+^] thereby ROS metabolism. **(A)** Volcano plot depicting whole-transcriptomic changes between indicated CD8^+^ T cell treatments. **(B)** Summary depiction of normalized enrichment scores (NES) of gene-sets with transcripts comparatively enriched or lost between indicated groups. **(C)** Representative GSEA comparisons of *Atp1a1* deficient vs sufficient CD8 T cells compared with MTORC1 Signaling and ROS associated datasets as indicated, summarized in **(B)**. **(D)** Summary depiction reactive oxygen species abundance (quantified by DCFDA fluorescence) and representative flow cytometry plots and summary data of indicated phenotypic surface proteins of CD8+ T cell populations treated with the antioxidant NAC. **(E)** Summary depictions of ROS as quantified by DCFDA and phenotypic surface markers quantified by flow cytometry of CD8+ T cells in the indicated conditions (↑ [K+]e = 40 mM). Error bars represent standard deviation. **P < 0.01, ***P < 0.001, ****P < 0.0001 (two-tailed t tests).

Extrapolating from these transcriptional changes, we found that *Atp1a1* deficient T cells harbor markedly elevated levels of ROS, quantified by the fluorometric dye DCFDA (**Figure 3D**). Recent observations that the antioxidant N-acetylcysteine (NAC) attenuates ROS, limits terminal differentiation and promotes stemness in T cells prompted us to apply NAC to *Atp1a1* deficient T cells to assess whether ROS accumulation was indeed the cause of their terminal differentiation (13). Antioxidant-mediated ROS neutralization prevented the accelerated differentiation otherwise observed following *Atp1a1* deletion, evidenced by normalization of PD-1, Tim-3, and other markers associated with advanced differentiation (**Figure 3D**). Moreover, isotonic K^+^ supplementation restored intracellular K^+^ levels (**Figure 1G**) and normalized intracellular ROS levels along with Fas, CD69, and CD62L levels (**Figure 3E**). In aggregate these findings indicate that intracellular [K^+^] itself prevents pathologic ROS accumulation and accelerated differentiation.

### The Na^+^-K^+^-ATPase is required for T cell antitumor activity in vivo

To assess the relevance of the Na^+^-K^+^-ATPase for T cell function *in vivo*, we adoptively transferred congenically distinguishable OT-I TCR transgenic CD8^+^ T cells following electroporation with Cas9-sgRNA complexes targeting *Atp1a1* or scramble sequences into C57BL/6 mice bearing established B16-OVA orthotopic melanoma. We found that *Atp1a1* deletion severely compromised T cell persistence after adoptive transfer within secondary lymphoid organs and the tumor site (**Figure 4A-C**). Demonstrative of their defect in persistence, both tumor regression and host survival were significantly impaired in mice receiving OT-I CD8^+^ T cells lacking *Atp1a1* (**Figure 4D,E**). Consistent with their compromised function, *Atp1a1* deficient OT-I CD8^+^ T cells were found in a terminally differentiated state within the tumor and spleen (**Figure 4F-I**). Collectively, these results indicate that the Na^+^/K^+^ ATPase plays a critical role in supporting T cell homeostasis and function by constraining pathologic accumulation of ROS and tonic signaling activity.

**Figure 4.**
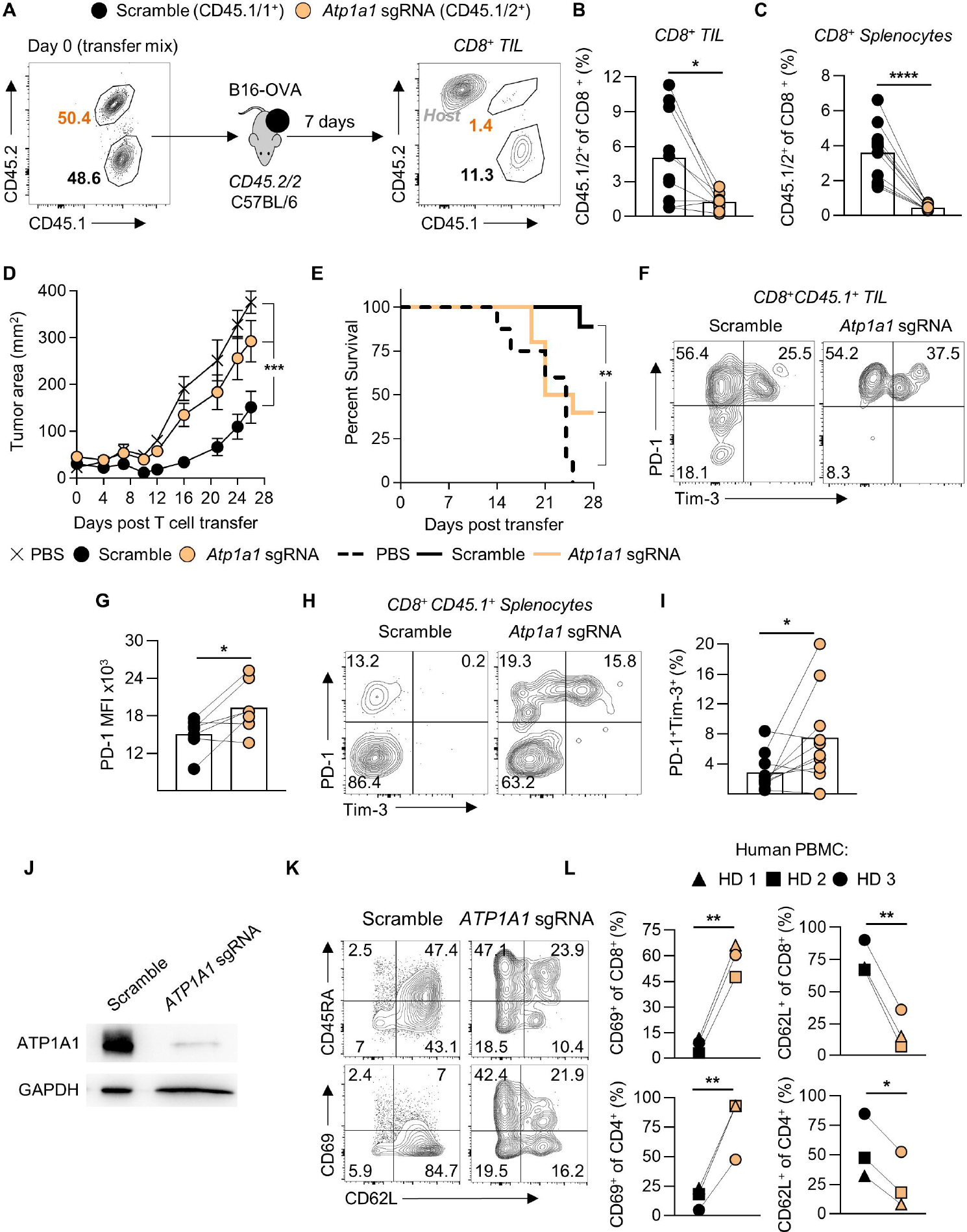
The Na^+^/K^+^ ATPase is required for cos+ T cell persistence and antitumor activity. **(A)** Pre-transfer flow cytometry analysis of OT-I CD8 T cells electroporated with Cas9 proteins complexed with scramble (CD45.1/1) or *Atp1a1* targeted (CD45.1/2) sgRNAs mixed at a ∼1:1 ratio prior to transfer to C57BL/6 (CD45.2/2) B16-OVA tumor bearing hosts. **(B)** Representative flow cytometry and summary analysis enumerating tumor infiltrating lymphocytes (TIL) and **(C)** spleen-resident transferred lymphocytes seven days after transfer to tumor-bearing mice as in **(A)**. **(D)** Rates of subcutaneous B16-OVA tumor growth represented over time following receipt of OT-I TCR transgenic CD8 T cells following Cas9 electroporation targeted with scramble or *Atp1a1* sgRNAs. **(E)** Kaplan-Meier survival estimate of the same cohort as in **(D)**. **(F)** Representative flow cytometry plot and summary data **(G)** depicting phenotypic state of OT-I CD8 T cells within B16-OVA tumors as in **(B)**. **(H)** Representative flow cytometry plot and summary data **(I)** depicting phenotypic state and enumeration of OT-I CD8 T cells within spleens of CD45.2/2 C57BL/6 mice bearing B16-OVA tumors as in **(B**,**C)**. **(J)** lmmunoblot quantification of ATP1A1 protein levels in primary human T cells derived from blood draws of healthy donors following electroporation with scramble sgRNAs or those targeting human ATP1A1 complexed to Cas9 proteins. **(K)** Representative flow cytometry plots of human CD8 T cells 5 days after electroporation with sgRNA-Cas9 complexes as in **(J)**, and **(L)** summary data of the same. Two-way ANOVA **(C);** ****P < 0.0001. Two-tailed Student’s t-tests **(B**, **C**, **F**, **H**, **K)**, *P < 0.05, **P < 0.01, ***P < 0.001. **B-D** and **E-H**, n=10 mice per group. Representative of two independent experiments. **I, J**, n=6-8 mice per group, data represents two pooled experiments.

Most importantly, Primary human CD8^+^ T cells obtained from multiple healthy donors demonstrated a similar requirement for intact Na^+^/K^+^ ATPase function. We confirmed disruption of ATP1A1 at the protein level via immunoblot (**Figure 4J**) and genomic via INDEL quantification (85-100%, data not shown). In all cases, genetic disruption of the Na^+^/K^+^ ATPase promoted the retention of the activation marker CD69 and loss of the memory associated lymphoid homing protein CD62L in both CD4^+^ and CD8^+^ T cells (**Figure 4 K,L**).

## Discussion

The gradients and transporters of monovalent ions have been primarily considered relevant for their role in supporting the electrical gradients required for ‘excitable’ cells (i.e. contain voltage-gated sodium channels) such as neurons, myocytes, or cardiomyocytes (14,15). While the importance of monovalent ion gradients in ‘non-excitable’ cells of hematopoietic origin has not been a topic of interest, prior work in T cells has hinged upon the importance of K_v_1.3 and K_Ca_3.1 in maintaining the electrical gradient (*V*_m_) to facilitate TCR induced Ca^2+^ influx (9,16,17). Building upon this work, we previously reported that isotonic elevations in extracellular K^+^ require PP2A phosphatase activity to limit Akt-mTOR signaling and effector function in T cells (1). Others have also confirmed that isotonic elevations to extracellular K^+^ do not impact TCR induced Ca^2+^ flux (18). Here, we again find that the impact of altering K^+^ transport in T cells cannot be accounted for by changes in Ca^2+^ alone.

While we did observe a depolarized resting membrane potential (*V*_m_) along with decreased Ca^2+^ influx following *Atp1a1* disruption, phenotypic and functional changes to T cell behavior otherwise were at odds with the anticipated impact of compromised Ca^2+^ influx. Rather than stem-like T cells with low NFAT activity, we found that T cells lacking *Atp1a1* harbored tonic phosphorylation of CD3ζ, among others, along with accelerated differentiation and formation of PD-1^+^Tim-3^+^ T_Ex_ cells. Transcriptomic assessment identified enrichment in the anabolic programs of c-Myc transcription along with mTOR metabolism despite an oxidative stress response. Following *Atp1a1* deletion, T cells harbored elevated levels of ROS and treatment with the antioxidant NAC restored ROS homeostasis and prevented terminal differentiation (PD-1^+^Tim-3^+^ T_Ex_ cells). Our observation that restoration of intracellular [K^+^] largely normalized ROS levels and T cell functional state underline the fundamental but largely unexplored role that intracellular [K^+^], independent of *V*_m_, plays in T cell function. Additionally, as ROS accumulation drove T_Ex_ cell formation, thereby compromising T cell antitumor activity *in vivo*, both ROS metabolism and its means of directing signal transduction are translationally relevant.

The term ‘ROS’ refers to a collection of molecules derived from oxygen and formed by reduction-oxidation reactions. The most prominent of these include superoxide (O_2_^-^), H_2_O_2_, along with organic hydroperoxides and radicals (19). Historically, ROS were considered harmful metabolic by-products that result from mitochondrial electron transport chain activity. More recently, ROS have been appreciated to also act as pleiotropic signaling co-factors derived from multiple sources that control both cellular homeostasis and response to external stimuli (20). Multiple reports have recognized the influence of ROS levels in T cells extends to TCR-induced signal transduction, nutrient consumption, and stemness (13,21,22). Relevant to both our prior and current findings, ROS accumulation results in the oxidation of phosphatase cysteine residues, suppressing their phosphatase activity (23). In this fashion, ROS-mediated phosphatase oxidation allows for effective kinase associated signal transduction by removing the inhibitory impact of ongoing phosphatase activity. ROS-induced cysteine oxidation has been observed in multiple phosphatases that are central to T cell function, such as PTP1B, SHP2, PP2A, and PTEN (19). In concert with our prior observations that K^+^-sensitive T cell function is enacted by phosphatase activity; these findings provide a framework in which intracellular K^+^ drives ROS abundance that subsequently influences the oxidation and function of relevant phosphatases. Taken together, these findings identify a previously unrecognized connection between ion transport, ROS homeostasis, and regulation of signal transduction in T cells that warrants further investigation from both a biological and translational perspective. Future studies will focus on how, specifically, intracellular K^+^ impacts ROS, determine the impact of ROS-independent consequences for T cell [K^+^] and transport, and assess inducible or contextual approaches to lower intracellular K^+^ transiently or mitigate the damaging consequences of elevated ROS. Broadly, these findings support a paradigm where T cell K^+^ transport can be used to change T cell behavior and improve cancer immunotherapies.

## Materials and Methods

### Study approval

All mouse experiments were performed in accordance to the guidelines issued by the Animal Care and Use Committee at OHSU using approved protocols. All infectious agents were approved by the OHSU Institutional Biosafety Committee.

### Mice and cell lines

C57BL/6 mice (obtained from Jackson Laboratories, Bar Harbor, ME) of 6–8 weeks of age were used as recipient hosts for adoptive transfer unless otherwise indicated. OT-1 Ly5.1 transgenic mice were used for adoptive cell transfer and viral kinetics experiments. To obtain OT-1 Ly5.1mice we crossed OT-1 (C57BL/6-Tg (TcraTcrb)1100Mjb/J) with Ly5.1 mice (B6.SJL-PtprcaPepcb/BoyJ) obtained from Jackson Laboratory. All mice were maintained under specific pathogen-free conditions. Modified B16-OVA, a mouse melanoma cell line, was transduced as previously described to express full-length ovalbumin protein in tandem with a T2A sequence and a blasticidin resistance insert; this line was used as the tumor model. Cell lines were maintained in complete media DMEM (Gibco) with 10% FBS, 1% glutamine and 1% penicillin–streptomycin.

### In vitro activation and culture of T cells

CD8+ T cells from OT-1 mice were stimulated in vitro with plate-bound anti-CD3 and anti-CD28 (5μg/ml; BioXcell) and expanded in RPMI 1640 containing 10% FBS and 100 IU/mL of recombinant human IL-2 (Peprotech). Human blood samples were enriched for T cells using negative bead selection (StemCell) and stimulated *in vitro* in media containing a 1:100 titer of TransAct (Miltenyi Biotec), 5% FBS and 100 IU/mL of rhIL-2. Cells treated with antioxidants were cultured with 10 mM N-acetylcysteine (Sigma). For measuring T cell effector, cells were washed of their original culture media (experimental or control and cytokines) prior to stimulation for 5 h with anti-CD3 and anti-CD28 without IL-2 in the presence of brefeldin A and monensin (BD Biosciences).

### Intracellular cytokine staining, phosflow and flow cytometry

For all flow cytometry experiments, T cells were stained with a fixable live/dead stain (Invitrogen) in PBS followed by surface antibody staining in FACS buffer (PBS with 0.5% BSA and 0.1% sodium azide). For intracellular cytokine staining, cells were first stained for surface markers and later stained for intracellular molecules following fixation and permeabilization following the manufacturer’s protocols (BD Cytofix/Cytoperm). For phosflow, T cells were incubated with αCD3 and αCD28-biotin conjugated antibodies (5 mcg mL^-1^) alongside live/dead and surface markers. TCR cross-linking was carried out at 37°C by the addition of soluble biotin (Thermo) to a final concentration of 20 µcg mL^-1^. Reactions were quenched at specified time point by the addition of warmed lyse/fix PhosFlow Buffer I (BD). Cells were then washed and permeabilized with – 80°C PhosFlow Buffer III (BD). After washing, cells were stained with phosphoantibodies purchased from Cell Signaling or BD Biosciences. Antibodies for surface staining and intracellular cytokine staining were purchased from BD Biosciences, BioLegend, eBiosciences and Invitrogen. All experiments were conducted on a BD (Becton Dickinson) Fortessa or Cytek Biosciences Aurora cytometer and analyzed with FlowJo software (Tree Star, Inc).

### Assessment of intracellular ROS, [K^+^]_i_ and *V*_m_

Cells were rested in IL-2-free complete RPMI 1640 for 16-18 hours. They were stained with fixable live/dead, surface markers, washed, then resuspended with 5 uM DiSBAC_2_(3) (Invitrogen), 2.5 uM DCFDA (AdipoGen Life Sciences) or 1 uM APG-4 (Asante Green-4, TEFLabs) in RPMI 1640 and incubated for 30, 20 and 90 minutes at RT, respectively. Cells stained for oxonol were washed 1X and analyzed by flow cytometry. DCFDA- and APG-4-loaded cells were analyzed on a cytometer immediately after incubation without a wash. For *V*_m_ assessment using DiSBAC_2_(3), to control for technical variations with dye loading, a fluorometrically distinguishable loading control was included within each sample and the quotient between the experimental cells and loading control cells is depicted as relative MFI. The ‘Relative MFI’ for both ‘Vehicle’ and ‘K^+^ 20 mM’ was determined using the average of ‘Vehicle’ replicates as the denominator. For determination of [K^+^]_i_, we generated standard curves representing induced [K^+^]_i_ vs. APG-4 MFI and used these standard curves to compute [K^+^]_i_ for each cell type. Briefly, standard curves accommodated [K^+^]_i_ ranging from 0 to 143 mM, where these [K^+^]_i_ were induced by incubating cells with isotonic, [K^+^]-controlled RPMI 1640, 1 uM APG-4 (pre-incubated 1:1 with Pluronic-127, Invitrogen) and K^+^-permeable ionophores valinomycin, gramicidin and nigericin at 10 uM. Cells queried for [K^+^]_i_ were incubated with APG-4 and isotonic, K^+^-free RPMI 1640. After flow analysis, we established linear regressions (r^2^ ≥ 0.80) and used them to deduce [K^+^]_i_ of the queried sample. Standard curves were generated for each unique cell type (i.e., *Atp1a1*-deficient cells, *Atp1a1*-sufficient cells).

### T cell receptor induced calcium influx

T cells were isolated and primed as above. Prior to analysis cells were rested in IL-2-free complete RPMI 1640 for at least 8 h. Cells were loaded with 1 μM Fluo3-AM and 1 μM Fura Red-AM (Invitrogen) for 30 min at 37 °C in HBSS with Ca2+, Mg2+ and 2% FCS, washed twice and then resuspended in HBSS with αCD3 and αCD28-biotin conjugates (eBioscience) and a live/dead stain (Invitrogen). For flow cytometry analysis, samples were resuspended in pre-warmed 37 °C additive-free RPMI, a baseline measurement was recorded for 20 s, followed by the addition of streptavidin (Invitrogen) to a final concentration of 20 μg mL^-1^ to induce TCR cross-linking and Ca^2+^ influx. Kinetic analyses were performed with the FlowJo software package (TreeStar).

### RNA sequencing and analysis

RNA-Seq and determination of differentially expressed genes was performed by Azenta Life Sciences (South Plainfield, NJ, USA). Briefly, RNA was extracted from frozen cell pellets using the Qiagen RNeasy Plus Universal Mini Kit per manufacturer’s instructions, then enriched for mRNAs using Oligod(T) beads. Following fragmentation for 15 mins at 94°C, cDNA was synthesized. cDNA fragments were end joined, adenylated at 3’ ends, adaptor ligated and enriched by PCR. Samples were clustered onto a flowcell and sequenced on the Illumina hiSeq 4000 (Illumina) with 2×150 bp paired end reads. Base-calling was executed by control software and raw sequence data were de-multiplexed using Illumina’s bcl2fastq 2.20 software. Sequenced reads were trimmed using Trimmomatic v.0.36 (USADEL LAB) and aligned to the mouse genome (ENSEMBL GRCm39) using the STAR aligner v.2.5.2b. Unique gene hit counts were deduced using the Subread package v.1.5.2, and only reads that fell within exon regions were retained for analysis. Differentially expressed genes were calculated using the R DESeq2 package. Fisher’s exact test or t-test were used to evaluate significance with indicated P value and fold-change thresholds. Gene set enrichment analysis was carried out using GSEA v4.2.3 software with enrichment of the MSigDB hallmark gene set (24,25) . RNA-Seq data will be deposited via GEO or provided as raw data for download.

### Immunoblot analysis

Western blot analysis was performed using standard protocols. In brief, for immunoblot quantifications, cells were resuspended in total cell extraction buffer and kept on ice for 10 min followed by homogenization. Cells were then centrifuged at 20,000g for 20 min at 4°C to pellet cell debris. Proteins were separated by 4%–12% SDS-PAGE, followed by standard immunoblot analysis using anti– Na^+^-K^+^-ATPase α1 or anti-GAPDH (Cell Signaling). Detection of proteins was performed using secondary antibodies conjugated to horseradish peroxidase-HRP and the super signal west pico chemiluminescent substrate (Thermo Scientific-Pierce).

### Statistical analysis

Data were analyzed using unpaired two-tailed Student’s *t* tests, two-way ANOVA. For adoptive transfer experiments, recipient mice were randomized before cell transfer. Tumor measurements were captured in a blinded fashion and plotted as the mean ± SEM for each data point, and tumor treatment graphs were compared by using the Wilcoxon rank sum test and analysis of animal survival was assessed using a log-rank test. In all cases, two tailed test with P values of less than 0.05 were considered significant. Statistics were calculated using GraphPad Prism 7 software (GraphPad Software Inc). Experimental sample sizes were chosen using power calculations, preliminary experiments, or were based on previous experience of variability in similar experiments. Samples which had undergone technical failure during processing were excluded from subsequent analysis.

### Retroviral transduction

Platinum-E ecotropic packaging cells (Cell Biolabs) were plated one day before transfections on poly-D-lysine-coated 10-cm plates (Corning) at a concentration of 6 × 10^6^ cells per plate. Packaging cells were transfected with 20 μg of retroviral plasmid DNA encoding MSGV-OVA-BSR, where indicated along with 6 μg pCL-Eco plasmid DNA using 60 μl Lipofectamine 2000 in OptiMEM (Invitrogen) for 8 h in antibiotic-free medium. Medium was replaced 8 h after transfection and cells were incubated for a further 48 h. Retroviral supernatants were then collected and spun at 2,000g for 2 h at 32 °C onto 24-well non-tissue-culture-treated plates coated overnight in Retronectin (Takara Bio) or spinfected for 90 minutes at 32°C in 1 µcg mL^-1^ polybrene, which was then removed after an additional 4 hour incubation.

### Adoptive cell transfer (ACT) and tumor immunotherapy

We implanted C57BL/6 were implanted with subcutaneous B16 melanoma (1 × 10^6^ cells). At the time of adoptive cell transfer (ACT), 10 days after tumor implantation, mice (n ≥ 5 for all groups) were sub-lethally irradiated (500 cGy), randomized, and injected intravenously with 2 × 10^6^ OT-1 Ly5.1 cells electroporated with scrambled sgRNA or Atp1a1-sgRNA. Mice used to assess day 7 post-transfer tumor infiltrating lymphocytes were intravenously injected with a mixture containing 6 × 10^5^ OT-1 (3 × 10^5^ of Scramble/ *Atp1a1* sgRNA treated each) cells total. Both cohorts received intraperitoneal injections of rhIL-2 in PBS (5 × 10^4^ IU in 0.5 mL) once daily for 2 days starting on the day of cell transfer. Tumors were measured using digital calipers. Tumor size was measured in a blinded fashion approximately every two days after transfer and tumor area was calculated as length × width of the tumor. Mice with tumors greater than 400 mm^2^ were euthanized. The products of the perpendicular tumor diameters are presented as mean ± s.e.m. at the indicated times after ACT.

### CRISPR-Cas9 mediated *Atp1a1* deletion and INDEL assessment

Non-targeting control sgRNA and *Atp1a1* targeted modified single guide RNAs (sgRNAs) were purchased from and synthesized by Synthego. *Atp1a1* loci were targeted with sgRNAs mixed in equal parts (murine *Atp1a1* sgRNAs AAAAGCCUCCAAAGAGCUGC, GAGGGGGUGUGAGGGCGUUG, CUUGCAGGUGUUAAUCCCUG; human ATP1A1 sgRNAs UGCUCGUGCAGCUGAGAUCC, CCAUUCAGGAGUAGUGGGAG, UGGAGCGAUUCUUUGUUUCU). Mixtures of complexed sgRNA and Cas9 (0.3 nmol synthetic sgRNA + 62 µmol Cas9 nuclease) and 2-10×10^6^ enriched murine CD8^+^ T cells were suspended in 25ul of P3 electroporation buffer (Lonza) and electroporated using the Lonza 4D Nucleofector (pulse code DN100), activated via 5 mcg/ mL anti-CD3 and anti-CD28 for 24 hours, and maintained in 100 IU mL^-1^ until day 4 of culture for analysis or adoptive transfer. Enriched human T cells were electroporated using pulse code EH115, activated using 1:100 TransAct and maintained in 200 IU mL^-1^ until analysis at day 5 of culture. For INDEL assessment, gDNA was harvested from cells at time of analysis and the region containing the *Atp1a1* sgRNA target was PCR amplified and Sanger sequenced. For murine samples, a 435-bp region was amplified for sequencing (forward primer: AGCAGAATGACCCAGAGTGG, reverse: GGGGGAGTTAACCAGGCTTC); for human samples, a 498-bp region was amplified (forward: GTGGGGACTGGCTCATCAG, reverse: GAGTTCATAACCATTAAGTAATGAGTGG). The indel percentage of amplicons derived from *Atp1a1-*deficient cells was deduced using the Synthego ICE tool (Synthego Performance Analysis, ICE Analysis. 2019. v3.0. Synthego).

## Acknowledgments

RE is supported by AACR MPM Oncology Charitable Foundation Transformative Cancer Research Grant, PanCAN Career Development Award, Conquer Cancer the ASCO Foundation Career Development Award, and NCI K08 CA256179.

